# Method to Identify Liver-to-Brain Region-Specific Crosstalk Using TurboID

**DOI:** 10.64898/2026.09.03.749208

**Authors:** Matthew C. Juber, Arvand Asghari, Kristin E. Claflin, Matthew J. Potthoff

## Abstract

Endocrine crosstalk between the liver and the brain regulates systemic energy homeostasis. However, this communication has not been comprehensively evaluated because identification of liver-derived proteins, or hepatokines, is difficult to define using bulk serum proteomics alone. To overcome this problem, we applied hepatocyte-restricted endoplasmic-reticulum–targeted TurboID (ER-TurboID) *in vivo* and recovered biotinylated, liver-secretion–competent proteins directly from specific brain regions, including the hypothalamus and hindbrain (nucleus of the solitary tract (NTS) and area postrema (AP)). Compared with AAV8-Null transduced controls, TurboID-expressing animals showed 3-to 21-fold enrichment of liver-expressed and predicted liver-secreted proteins among brain regions, with 81 canonical liver-secretome proteins, including multiple carboxylesterase (CES) family members. These results establish a method to identify novel secreted factors from peripheral tissues that can potentially act on specific brain regions.

## Introduction

The liver is a central node of inter-organ communication, releasing a diverse secretome consisting of complement and coagulation factors, apolipoproteins, serpins and acute-phase proteins, carrier proteins, hepatokines, and detoxification enzymes such as the carboxylesterase (CES) family. The production of these factors coordinates systemic responses to nutritional, environmental, and metabolic stress [1]. A subset of these proteins acts directly on the central nervous system to regulate energy homeostasis. For example, the liver-derived hormone fibroblast growth factor 21 (FGF21) signals to the hypothalamus and hindbrain to modulate energy expenditure, food preference, and sympathetic outflow [2, 3]. By identifying the liver-derived proteins which signal to the brain, new modes of peripheral to central nervous system (CNS) communication may be revealed which could define new pathways regulating systemic metabolic homeostasis.

Previous attempts to characterize the liver secretome have been constrained by multiple technical hurdles. Performing bulk plasma proteomics catalogues circulating proteins but cannot determine the source or destination of identified factors. In addition, there are technical hurdles in identifying lowly expressed proteins like hormones because of the extensive quantity of other proteins in circulation. While gene expression profiling could be used to identify factors that are predicted to be secreted, tissue-specific changes in secretome production could change rapidly without changes in gene expression (e.g., insulin release). In vivo proximity labeling with the engineered biotin ligase TurboID [4] has recently been used to tag secreted proteins as they are synthesized in defined cell types in vivo [5-7]. Interestingly, this strategy has identified novel exercise-induced hepatokines from plasma [7]. However, plasma-based recovery still does not demonstrate that a biotinylated, liver-secreted protein reaches a specific target tissue. This methodological gap is particularly limiting for liver-to-brain communication.

The hypothalamus and hindbrain are two critical brain regions that integrate peripheral nutrient, hormonal, and gut-derived signals to control energy and nutrient homeostasis [8-10]. Hypothalamic neurons are responsive to glucose, amino acids, fatty acids, insulin, and leptin [11], and the hindbrain, including the nucleus of the solitary tract (NTS) and area postrema (AP), receive input from gut-derived nutrient signals and circulating leptin and ghrelin [12, 13]. However, the signals that are sensed by these brain regions may not be complete and have not been comprehensively evaluated.

To begin to address this issue, we developed an *in vivo* strategy that traces liver-derived proteins to defined brain regions by combining hepatocyte-restricted, ER-targeted TurboID expression with streptavidin enrichment and mass spectrometry of dissected hypothalamus and NTS/AP. Here we show that biotinylated, liver-secretion–competent proteins are recoverable from both brain regions and are several-fold enriched for liver-expressed and predicted-secreted proteins relative to AAV8-Null controls. We further demonstrate that 81 canonical hepatic-secretome proteins spanning every major functional class — complement, coagulation, apolipoproteins, serpins, carrier proteins, hepatokines, and the CES family — are significantly enriched in the brain in TurboID-expressing animals. Notably, multiple CES family members, including Ces2a and Ces2c, are consistently recovered in both the hypothalamus and NTS/AP, identifying the brain as a candidate target tissue for liver-secreted CES proteins. Together, these results establish hepatocyte-restricted in vivo proximity labeling as a direct, spatially resolved approach for mapping hepatic protein delivery to the CNS.

## Material and methods

### Animals and viral Constructs

TBG-ER-TurboID was obtained from Addgene (149415) and produced on an AAV8 vector by VectorBuilder. AAV8-TBG.PI.Null.bGH was procured from Addgene (105536-AAV8) for an empty vector control. 8–10-week-old C57BL6/J anesthetized mice were transduced retro-orbitally with 10 µL of AAV8-TBG-ER-TurboID or AAV8-TBG.PI.Null.bGH diluted in 100 µL PBS. Mice were given 1 week to recover from injections prior to experiments. Mice were individually housed in a 12 h light/dark cycle at 22-23°C and were given *ad libitum* access to chow (Teklad; 2920X). Health status was normal for all animals. All experiments presented in this study were conducted according to the animal research guidelines from NIH and were approved by the University of Iowa IACUC.

### Biotin administration

Mice were given access to water supplemented with 50 mg/mL biotin (Sigma B4501) 1 week after viral transduction for 5 days. Biotin-water solution was adjusted to a neutral pH (7-7.5). On the 5^th^ day of biotin-water access, mice were anesthetized and perfused with 30 mL ice-cold PBS after serum collection to reduce or clear biotinylated proteins and non-specific binding to the microvasculature. Tissues were harvested and immediately snap frozen.

### Immunoprecipitation of biotinylated proteins

Individual brain regions were pooled such that tissue from five animals was combined to generate one representative sample, yielding n = 3 pooled samples per group per brain region (15 mice per group total). Immunoprecipitation was performed as previously described [7] adapted for tissue lysate enrichment. In brief, pooled tissue was homogenized in cold RIPA buffer using MP Biomedicals Bead Beater homogenizer at 4 °C and cleared by centrifugation (13,000 rpm, 10 min, 4 °C), and protein concentration was determined by BCA. Free biotin was removed from the lysates with 3 kDa centrifugal filter units (MilliporeSigma,UFC900308) prior to incubation with streptavidin beads. Dynabeads MyOne Streptavidin T1 magnetic beads were washed twice in washing buffer (50 mM Tris-HCl, 150 mM NaCl, 0.1% SDS, 0.5% sodium deoxycholate, 1% NP-40, 1 mM EDTA, 1× HALT protease inhibitor, 5 mM Trolox, 10 mM sodium azide, 10 mM sodium ascorbate) and combined with lysate at 1 µl beads per 10 µg protein. Beads were incubated overnight at 4 °C with rotation. Beads were then washed twice with 1 ml RIPA buffer, once each with 1 ml 1 M KCl, 1 ml 0.1 M Na_2_CO_3_, and 1 ml 2 M urea in 10 mM Tris-HCl (pH 8.0), and twice with 1 ml washing buffer. Bound proteins were eluted by boiling at 95 °C for 10 min in 60 µl 2× sample buffer supplemented with 20 mM DTT and 2 mM biotin. Eluates were stored at −80 °C until processing for proteomics.

### Western Blot Analysis

Western blot analysis was performed as previously [14]. Briefly, snap-frozen tissues were homogenized in RIPA buffer with protease and phosphatase inhibitors. Samples were centrifuged at 13,000xg for 10 minutes and infranatant was collected. Protein content was quantified using BCA assay. Samples were diluted to a standard concentration, mixed with an appropriate amount of Laemmli buffer, and incubated for 10 minutes at 100°C, then stored at -20°C. Equal protein amounts were loaded for SDS-PAGE on 10% acrylamide gels. Proteins were transferred to a PVDF membrane via dry transfer (BioRad; Trans-Blot Turbo) then probed with the specified primary antibodies. The following primary antibodies were used: anti-V5 (Invitrogen, R960-25) and anti-biotin (Invitrogen, S21378). No-Stain Protein Labeling Reagent (Invitrogen, A44449) was used to examine total protein loading.

### Proteomics Sample Prep and LC-MS/MS Analysis

Label-free quantitative proteomics was performed by the Duke Proteomics and Metabolomics Core Facility. Streptavidin-enriched protein samples from hypothalamus and NTS/AP were processed using S-Trap micro cartridges following reduction, alkylation, and spiking with internal QC standards. Proteins were digested with trypsin and peptides were eluted, lyophilized, and resuspended prior to analysis. Peptides were analyzed on a Thermo Orbitrap Astral mass spectrometer coupled to an Evosep One UPLC system using a data-independent acquisition (DIA) method. Data was searched against the Mus musculus SwissProt database using a library-free approach in Spectronaut. Peptide intensities were filtered, normalized by total signal intensity and a trimmed mean method, then aggregated to the protein level for quantification. Reproducibility was confirmed by low %CV in pooled QC and replicate groups, and differential expression was assessed by fold change and heteroscedastic t-tests on log_2_-transformed data.

### Bioinformatic Analysis

Differential abundance was assessed using limma moderated t-statistics (eBayes with trend = TRUE) on log_2_-transformed Spectronaut-normalized intensities, with Benjamini–Hochberg adjustment to control the false discovery rate. Proteins were considered significantly enriched when |log_2_ fold change| ≥ 1.5 and adjusted p-value (q) < 0.05.

To define a panel of bona fide hepatic-secretome candidates, we curated a list of canonical liver-secreted gene symbols across nine functional classes (complement components; coagulation factors and pathway-related serpins; apolipoproteins; other serpins and acute-phase proteins; pattern-recognition and innate-immunity proteins; carrier proteins; hepatokines; the carboxylesterase family; and the hepatic IGF-1–axis hormone receptor Ghr), compiled from textbook hepatology and the MDSEC reference. Proteins matching this panel that also met the limma significance threshold (|log_2_FC| ≥ 1.5, q < 0.05) in the TurboID-vs-Null comparison in at least one brain region were classified as bona fide hepatic-secretome hits (n = 81; Fig. 2).

To score liver origin, we integrated three independent positive sources of evidence: UniProt [15], the Human Protein Atlas [16], and a curated mouse hepatocyte-secretome whitelist. UniProt tissue-specificity text was parsed for positive mentions of liver or hepatocytes at the sentence level, rejecting negation contexts (e.g., “not in liver”). HPA entries categorized as Tissue enriched, Tissue enhanced, or Group enriched with liver listed in the RNA tissue-specific nTPM field were flagged as liver-enriched. The curated whitelist (162 genes) supplemented mouse-specific paralogs that human-centric resources miss, including the Serpina1 family, the major urinary protein family, mouse SAA paralogs, and select complement and coagulation factors. Proteins satisfying any of the three criteria were flagged Liver-expressed; those additionally lacking a UniProt brain mention were Liver, non-brain; combination with a UniProt-predicted signal peptide defined the highest-confidence Liver-secreted (predicted) subset (Figs. 1C, 1E).

**Figure 1.**
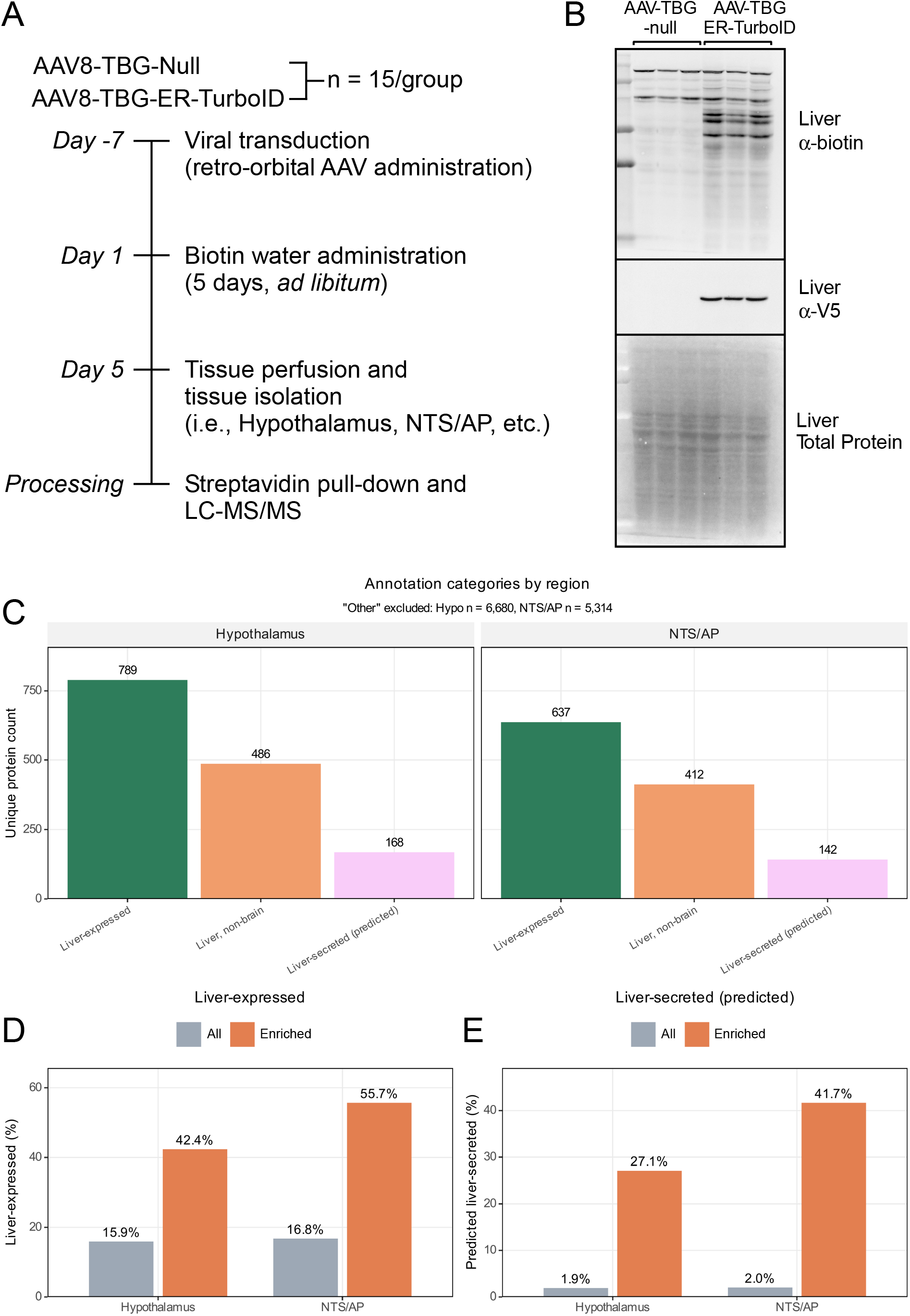
Experimental Design and Protein Annotation Overview. (A) Schematic of the *in vivo* ER-TurboID labeling experiment. 8–10-week-old wild-type male C57Bl/6J mice on chow diet were transduced with AAV8-TBG-Null or AAV8-TBG-ER-TurboID via retro-orbital injection (n = 15/group). One-week post-injection, mice received biotin-supplemented water for five days to maximize hepatic TurboID labeling. Hypothalamus and nucleus of the solitary tract, area postrema (NTS/AP) tissues were harvested from each mouse, and then individual brain regions were pooled such that tissue from five animals was combined to generate one representative sample, yielding n = 3 pooled samples per group per brain region (i.e., 5 hypothalami were combined together for one sample three times (for a total of n=3), and the same was performed separately for the NTS/AP). Streptavidin-enriched lysates were then subjected to LC-MS/MS proteomic analysis. (B) Western blot analysis for ER-TurboID (V5-tagged) and hepatic biotinylated proteins in 8–10-week-old wild-type male C57Bl/6J mice on chow diet transduced with AAV8-TBG-Null or AAV8-TBG-ER-TurboID via retro-orbital injection. (C) Number of unique proteins detected per brain region (hypothalamus and NTS/AP) by annotation category (Liver-expressed; Liver, non-brain; Liver-secreted (liver-expressed + not expressed in brain + signal peptide). Two non-informative categories are excluded for clarity: “Other” (Hypothalamus n = 6,680; NTS/AP n = 5,314) and “Signal Peptide Only” (Hypothalamus n = 932; NTS/AP n = 591). (D) Percentage of detected proteins annotated as liver-expressed among all detected proteins versus those significantly enriched by limma (|log_2_FC| ≥ 1.5, q < 0.05). Liver-expressed enrichment is 2.7-fold in hypothalamus (15.9% → 42.4%) and 3.3-fold in NTS/AP (16.8% → 55.7%). (E) Same comparison for the highest-confidence subset (liver-expressed, brain-excluded, with predicted signal peptide; Liver-secreted). Enrichment is 14.6-fold in hypothalamus (1.9% → 27.1%) and 20.8-fold in NTS/AP (2.0% → 41.7%).

**Figure 2.**
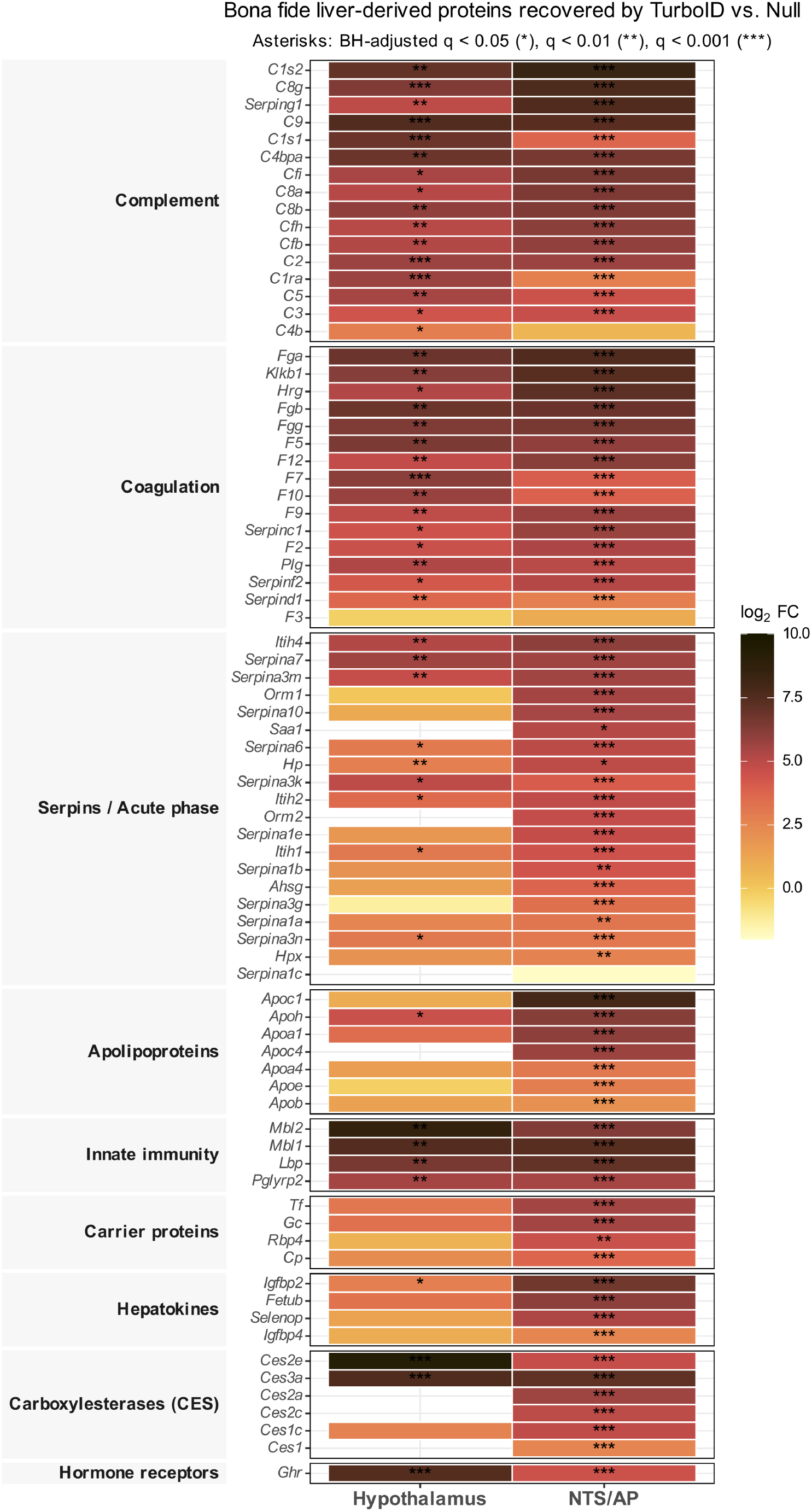
Eighty-one canonical liver-derived proteins, spanning every major class of the hepatic secretome, are recovered from the hypothalamus and NTS/AP. Heatmap of all proteins from the curated bona fide hepatic-secretome list (n = 81 across nine functional classes) that met the significance threshold in the TurboID-vs-Null comparison in either the hypothalamus or nucleus of the solitary tract, area postrema (NTS/AP) brain regions. Color: limma moderated log_2_ fold change (lajolla palette; deeper red indicates stronger enrichment). Asterisks indicate Benjamini-Hochberg (BH)-adjusted significance: * q < 0.05; ** q < 0.01; *** q < 0.001. Rows are grouped by functional class (Complement, Coagulation, Serpins / Acute phase, Apolipoproteins, Innate immunity, Carrier proteins, Hepatokines, Carboxylesterases, Hormone receptors) and ordered within class by maximum log_2_ fold change.

## Results

### Liver-Expressed Protein Detection in Brain Regions Following TurboID Labeling

To identify liver-derived proteins capable of reaching the brain, we employed a proximity biotinylation approach using TurboID targeted to hepatocytes (Fig. 1A). Mice were transduced with AAV8-TBG-ER-TurboID to drive ER-TurboID expression restricted to liver, followed by five days of biotin administration via drinking water (Fig. 1A). AAV-TBG-ER-TurboID administration resulted in expression of ER-TurboID in the liver and increased protein biotinylation (Fig. 1B), as expected. Mice were then perfused with excess cold PBS to remove circulating proteins from circulation and tissue vasculature as much as possible.

Mass spectrometry of proteins from pooled streptavidin-purified hypothalamus and nucleus of the solitary tract, area postrema (NTS/AP) samples revealed a wide array of proteins, which we annotated for liver expression, brain expression, and signal peptide presence. Proteins were classified into three informative categories: (1) Liver-expressed, (2) Liver expressed, but not expressed in brain (liver, non-brain), and (3) Liver-secreted (expressed in the liver, not expressed in the brain, and predicted to be secreted) (Fig. 1C). From the hypothalamus, 789 proteins were liver-expressed, 486 were liver-expressed and not expressed in brain tissue, and 168 met all three criteria (liver-expressed + not expressed in brain + signal peptide). Similar trends were observed in the NTS/AP, albeit with fewer total proteins: 637 were liver-expressed, 412 were liver-expressed and not expressed in brain, and 142 met all three criteria (liver-expressed + not expressed in brain + signal peptide). These annotations demonstrate that a significant subset of proteins detected in both brain regions are consistent with hepatic origin and secretory potential, supporting the feasibility of using ER-TurboID for tracing liver-to-brain protein communication.

### Significantly Enriched Liver-Derived and Secreted Proteins

To determine whether proteins significantly enriched in the brain following liver-targeted TurboID labeling were enriched for liver-derived secretory proteins, we compared annotation distributions between all detected proteins and those significantly enriched by limma moderated t-statistics (|log_2_FC| ≥ 1.5, q < 0.05) in the hypothalamus and NTS/AP. In both regions, significantly enriched proteins were more frequently annotated as liver-expressed than in the background. In the hypothalamus, 42.4% of enriched proteins were liver-expressed versus 15.9% of all detected proteins; in the NTS/AP, the enrichment was greater (55.7% vs. 16.8%) (Fig. 1D).

Restricting to the highest-confidence subset — liver-expressed, brain-excluded, with a predicted signal peptide (Liver-secreted (predicted)) — the enrichment became more dramatic. In the hypothalamus, 27.1% of enriched proteins were predicted liver-secreted versus 1.9% of all detected proteins (14.6-fold), and in the NTS/AP, 41.7% versus 2.0% (20.8-fold) (Fig. 1E). Across the ER-TurboID-vs-Null comparisons in hypothalamus and NTS/AP, 81 individual proteins of canonical hepatic origin met our significance threshold, spanning every major class of the hepatic secretome: complement components (C1s1/2, C2, C3, C5, C8a–g, C9, Cfh, Cfi, Cfb, Serping1), coagulation factors (F2, F5, F7, F9, F10, F12, Fga, Fgb, Fgg, Plg, Klkb1, Hrg), apolipoproteins (Apoa1, Apoa4, Apob, Apoc1, Apoc4, Apoe, Apoh), serpins and acute-phase proteins (Serpina1a– e, Serpina3g/k/m/n, Serpina7, Hp, Orm1, Orm2, Itih1/2/4), pattern-recognition and innate-immunity proteins (Mbl1, Mbl2, Lbp, Pglyrp2), carrier proteins (Tf, Cp, Gc, Rbp4, Ahsg), hepatokines (Selenop, Igfbp2, Igfbp4, Fetub), and the carboxylesterase (CES) family (Fig. 2). These data show that liver-specific, secreted proteins are recoverable from the brain when biotinylated and streptavidin-purified.

## Discussion

This work introduces a method for mapping inter-organ protein communication by combining tissue-restricted TurboID expression (signal sender) with recovery of biotinylated proteins from a separate tissue not expressing TurboID (signal receiver). Unlike prior approaches that profile secretomes from plasma or conditioned media, this strategy enables direct detection of biotinylated proteins in target organs, providing spatial resolution of protein trafficking.

Enriching biotinylated proteins from complex tissues like the brain introduces technical caveats. Streptavidin-based purification, while highly efficient, can co-purify non-biotinylated proteins through nonspecific binding or association with endogenous biotin-containing proteins. To address this, we used AAV8-Null transduced animals to establish a per-region detection background and applied limma moderated t-statistics with BH-adjusted q-values to identify proteins differentially enriched above that background. Candidates were further filtered for liver origin (UniProt tissue specificity, Human Protein Atlas tissue enrichment, and a curated mouse hepatocyte-secretome whitelist) and predicted secretion (UniProt signal peptide).

Using this strategy, we identified canonical liver-derived proteins spanning complement, coagulation, apolipoprotein, serpin, carrier, and hepatokine classes — including multiple CES family members — consistently significantly enriched in both the hypothalamus and NTS/AP. Among the recovered proteins, the carboxylesterase (CES) family is of particular interest. CES proteins are involved in phase-I drug metabolism, lipid metabolism, and detoxification of environmental and endogenous toxins [17]. CES proteins are less commonly reported as circulating factors, yet six CES family members (Ces1, Ces1c, Ces2a, Ces2c, Ces2e, Ces3a) were consistently recovered in NTS/AP. A previous report discovered CES2 as being secreted and functioning to improve metabolic health [7], and our data confirms that CES2 is not only secreted but also suggests that it may target brain regions to mediate some of its systemic effects.

In addition to the technical caveats, there are several additional limitations to the ER-TurboID approach we employed to identify tissue crosstalk. While we know the source of the signals (i.e., the liver), our method cannot determine which cell types within the brain regions (i.e., neurons, glia, or resident immune cells) are the exact targets of the liver-derived proteins. In addition, it is unclear whether these signals are actually biologically active. Another issue is that some proteins had incomplete annotations in UniProt and HPA, which could underestimate the breadth of recovered liver-derived candidates. There is also intrinsic risk of nonspecific binding with streptavidin enrichment, and proteins in complex with endogenous biotin can contribute additional false positives. Finally, as we used 3 kDa filter tubes to remove free biotin, this method would not identify any proteins smaller than 3 kDa. Nevertheless, this method provides an important advance in the identification of novel tissue crosstalk.

## Acknowledgements

This work is funded by the National Institutes of Health (NIH) R01DK106104 (M.J.P.), Veterans Affairs Merit Review program I01BX004634 (M.J.P.), and K01DK133667 (K.E.C.). The authors would like to acknowledge the use of Duke University Proteomics and MetabolomicsCore Facility.

